# The role of amygdala calcitonin gene-related peptide receptors on the development of persistent bladder pain in mice

**DOI:** 10.1101/2025.06.10.658965

**Authors:** Lakeisha A. Lewter, Blesson K Paul, Arnold M. Salazar, Uma Chatterjee, Hoai Phuong T Pham, Myra Z. Khan, Anna E. Schmitz, Abraham M. Nofal, Mursal M. Hussein, Indira U. Mysorekar, Benedict J. Kolber

**Author notes:** Corresponding Author: Benedict Kolber, University of Texas at Dallas, Richardson, TX.

## Abstract

Bladder pain significantly impacts millions worldwide, severely affecting their quality of life and posing a major clinical challenge. Understanding the mechanisms underlying persistent bladder pain is critical for developing better therapeutic strategies. In this study, we investigate the effects of cyclophosphamide (CYP)-induced persistent bladder sensitization to explore the lateralized contribution of amygdala calcitonin gene-related peptide receptors (CGRP-Rs) on pain-like changes in mice. We demonstrate that CYP induces hypersensitivity lasting up to 14 days post-injury (DPI) in the urinary bladder distention assay and up to 21 DPI when assessing abdominal mechanical sensitivity. Despite persistent pain-like changes, no significant bladder histological changes were observed. Based on previous findings that CGRP signaling from the parabrachial nucleus contributes to central amygdala (CeA) lateralization, we hypothesized that CGRP-Rs play a key role in driving visceral bladder pain-related hemispherical differences. We show that inhibiting CGRP-R activity with the antagonist CGRP_8-37,_ in the right CeA attenuates bladder pain-like behavior, whereas left CeA inhibition sustains CYP-induced hypersensitivity. Electrophysiological recordings revealed increased firing frequency in CGRP-R positive cells in the right CeA 7 DPI. *In vivo* single photon calcium imaging demonstrated increased Ca transients in CGRP-R-positive cells in the right CeA, upon the presentation of a stimulus at 0 DPI, and overall at 2DPI, further confirming the pronociceptive role of CGRP-Rs in the right CeA. Taken together, these findings provide a crucial foundation for understanding pain-induced CeA lateralization and for further studies identifying how targeting CGRP signaling could provide bladder pain relief.

## Introduction

While the majority of preclinical pain research focuses on somatic pain, visceral pain, such as interstitial cystitis/bladder pain syndrome (IC/BPS), remains understudied^1^. IC/BPS is a long-term, highly distressing condition that affects the urinary bladder^2^. It is typically marked by frequent and urgent urination, ongoing abdominal and pelvic pain^2^. IC/BPS is categorized under the umbrella term urologic chronic pelvic pain syndrome (UCPPS). UCPPS affects more than 10 million people in the United States^3^. Research has suggested several possible contributors to UCPPS such as defective urothelial integrity and function, changes in sensitization and neuroplasticity, infectious events, and inflammation^4^. Despite these research efforts, the etiology of bladder pain and other pelvic pain conditions is still poorly understood. Understanding the underlying mechanisms involved in bladder pain progression from acute to chronic is essential for developing novel and effective therapeutic interventions.

IC/BPS is often comorbid with affective disorders such as depression and anxiety, suggesting that the central nervous system plays a role in bladder pain processing^5^. The amygdala has been identified as a key component of the pain matrix^6^. Anatomical, behavioral, and physiological changes have been observed within the amygdala in various injury states^6–10^. In particular, a subdivision of the amygdala, the central nucleus of the amygdala (CeA), has been shown to be a major site of nociceptive processing, receiving input through the spino-parabrachio-amygdaloid pathway or through a thalamic-cortical-basolateral nucleus of the amygdala (BLA) relay^11^. Additionally, there are two other characteristics of the CeA that make this particular brain region worth exploring in the context of pain. First, hemispheric left-right differences of the CeA have been observed in the context of pain. Both animal and human studies show evidence of amygdala lateralization in the context of pain^12,13^. In human studies, an increase in regional gray matter was observed in the left amygdala in UCPPS patients, compared to healthy patients or those with other visceral pain conditions^14^. In animal studies, the role of the right CeA has been shown to be dominant in producing pronociceptive effects^7,8,15–17^. The left CeA has been shown to have diverse and sometimes contrasting effects in models of somatic pain, but has shown antinociceptive functions in mouse models of visceral bladder pain^18,19^.

Second, the CeA is heterogenous, consisting of cells expressing a variety of neuropeptides and neuropeptide receptors. Previous work in our lab has shown that when calcitonin gene-related peptide (CGRP) is injected into the right CeA, animals display an increase in bladder pain-like behavior, whereas a decrease in bladder pain-like behavior is observed when CGRP is injected in the left^19^. These data suggest that CGRP contributes to amygdala lateralization in the context of bladder pain-like changes.

What is not understood is how CeA lateralization changes as bladder pain transitions beyond the immediate acute state. In a rat neuropathic pain model, spontaneous and stimulus-evoked neuronal activity in the left CeA was increased at 2 and 6 days after neuropathic injury, but lateralization transitioned to an increase in neuronal activity in the right CeA 14 days after injury^20^. 14 days after injury corresponds to other markers of chronic pain^21^, suggesting that the development of right CeA activity may contribute to the chronic pain transition. It is unknown whether this shift of dominant neuronal activity from the left to the right CeA is also present in visceral bladder pain. Furthermore, the role of CGRP receptors (*Calcrl* or CGRP-Rs) on the transition from acute to chronic/persistent bladder pain has not yet been explored. We hypothesized that CGRP-Rs play a key role in driving visceral bladder pain-related lateralization of the amygdala. We predict that upon injury, the antinociceptive effect of CGRP is gradually lost in the left CeA, while the pronociceptive effect of CGRP is gradually enhanced in the right CeA.

In this study, we tested this hypothesis using behavioral pharmacology, evaluation of bladder histology, slice electrophysiology, and *in vivo* calcium imaging, to determine the influence of amygdala CGRP receptors on the development of persistent bladder pain across hemispheres.

## Materials and Methods

### Animals

Experiments used adult (8-13 weeks) male and female C57BL/6J (Jackson Laboratory stock #000664), Calcrl^tm1.1(cre)Rpa^ (*Calcrl*^Cre^)(kindly provided by Dr. Richard Palmiter, University of Washington^22^) and Ai14(RCL-tdT)-D (Jackson Laboratory stock #007914) mice^23^. C57Bl/6J mice were used for bladder physiology, histology, and behavioral analysis in wildtype mice. *Calcrl*^Cre/wt^ mice were generated from *Calcrl*^Cre/Cre^ x C57Bl/6J mice and were used for single photon calcium imaging as described below. *Calcrl*^Cre/wt^ x Ai14(RCL-tdT)-D^+/-^ mice were mated to generate heterozygotes of each genotype. These mice were used to mark *Calcrl* positive cells during *ex vivo* slice physiology experiments as described below. Details on each line is described below related to individual experiments. All mice were backcrossed to the C57BL/6J strain.

Mice were group-housed (except after stereotaxic surgery) in a controlled environment with a 12-hour:12-hour light:dark cycle in the vivarium. Food and water were made available *ab libitum* except during experimental sessions. For wildtype mice purchased from Jackson Laboratory, experiments began once animals were acclimated to the University’s Animal Facility for at least 72 h. Male and female mice were used for all experiments unless otherwise noted. Animals were maintained and experiments were approved and conducted in accordance with the guidelines established by the Institutional Animal Care and Use Committee (IACUC) at The University of Texas at Dallas (protocol 20-04 and protocol 2023-0092).

### Cyclophosphamide Bladder Sensitization

Cyclophosphamide (CYP; Sigma-Aldrich cat # C0768) dissolved in sodium chloride 0.9% normal saline was used to induce bladder pain-like sensitivity in mice^24,25^. Mice received 100 mg/kg CYP (intraperitoneally) every other day for 5 consecutive days (3 total injections). Behaviors were tested before and after (1-21 days post-injury (DPI)) CYP treatment. 24 hours after the first injection of CYP, is considered 1 day post-injury (1 DPI).

### Urinary Bladder Distension – Visceromotor Response (UBD-VMR)

The sample size used in this experiment was based on a previous UBD-VMR study in bladder-sensitized mice^19^. Female C57BL/6J mice were placed under 2% isoflurane and underwent surgery to expose the external oblique abdominal muscle, as briefly described^26^. Two electrodes were placed in the left external oblique muscles, while a grounding electrode was attached near the chest region. We previously demonstrated that the side of the body recorded does not impact brain lateralization^18^. A lubricated 24-gauge, 14mm catheter was inserted into the bladder via the urethra. Mice were catheterized and the isoflurane was gradually decreased in 0.125% increments every 10 minutes until 0.8% isoflurane was reached and body temperature was maintained at 37°C throughout consistent with our established methods^27^. Electromyogram (EMG) signals were recorded through a P511 Grass Amplifier connected to CED 1401 ADC and Spike 2 software (CED, v. 10.07). Once electromyography responses were stable, pressurized air was delivered via a catheter at different pressures (30 mmHg and 60 mmHg) using a custom visceral pressure stimulator^28^. Data were exported to Matlab where the background EMG was subtracted and the stimulus evoked EMGs were rectified and integrated over the 20 second pressure period using a custom Matlab script. For UBD-VMR, mice were randomly assigned to 4 different groups. One group was treated with saline, and 4 groups were treated with CYP. The saline group is referred to as 0 DPI group throughout this study. However, saline was given 1-2 days prior to the UBD-VMR test. For the CYP-treated groups: the first injection of CYP was given 2 days (2 DPI), 7 days (7 DPI), 14 days (14 DPI), and 21 days (21 DPI) prior to testing (**Fig. 1A**).

**Figure 1:**
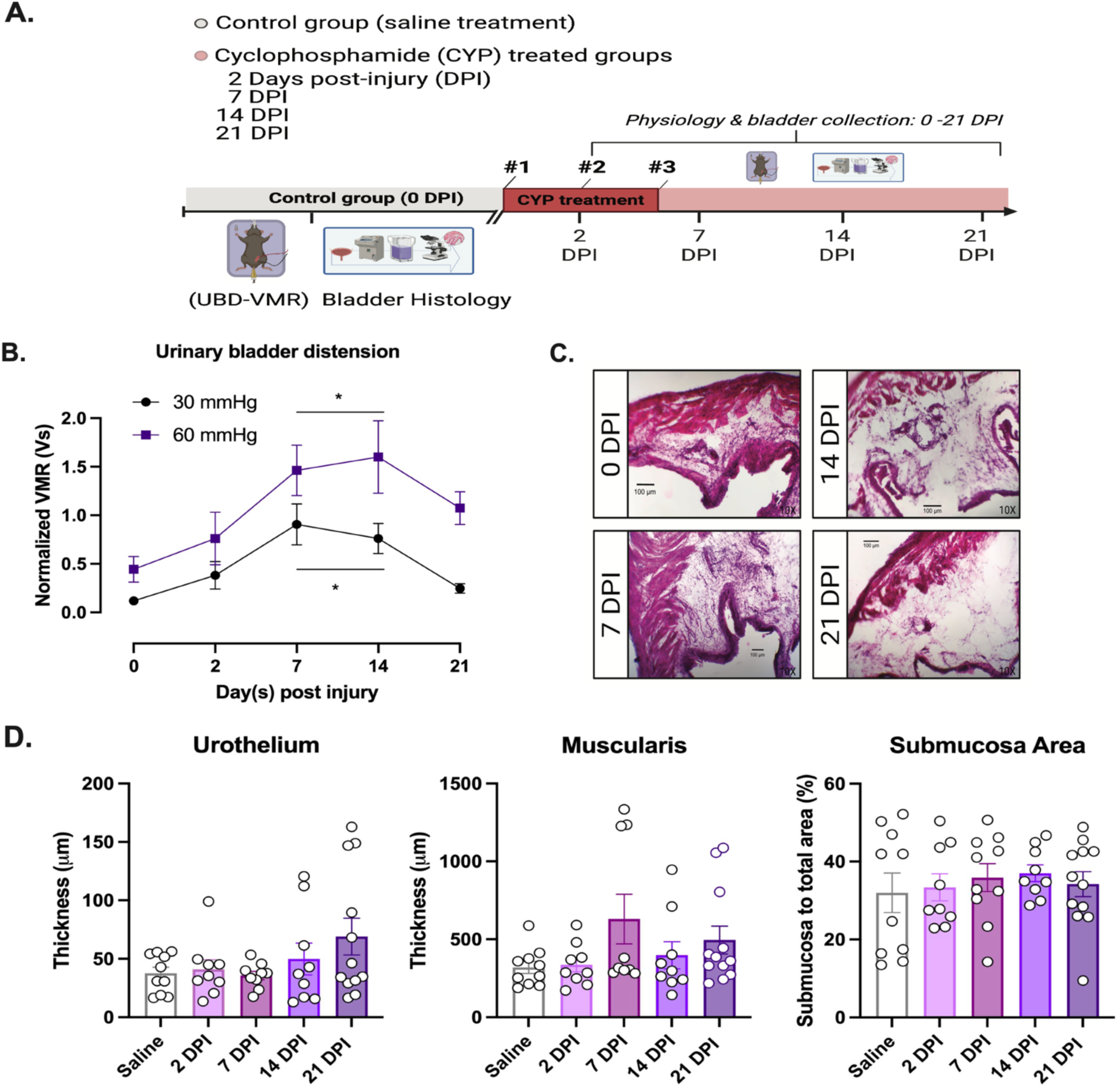
Cyclophosphamide increases hypersensitivity 7 and 14 days post injury, without disrupting bladder histology. **A)** Schematic of experimental timeline. **B)** Urinary bladder distension – visceromotor responses (UBD-VMR) in saline-treated (0 DPI) and CYP-treated (2-21 DPI) female mice at innocuous (30 mmHg-black circles) and noxious (60 mmHg-purple squares) pressures (n=8-10/group). Asterisks represent results from Tukey’s posthoc test. Tukey’s multiple comparisons test reveals a significant increase at 7 and 14 DPI (p< 0.05, *). **C)** Representative images of bladder sections after hematoxylin and eosin staining at 0, 7, 14, and 21 DPI. **D)** Quantification of the urothelium, muscularis, and submucosa area of bladders from saline-treated and CYP-treated mice at different timepoints (n=8-10/group). All results are presented as mean ± SEM.

### Hematoxylin and Eosin (H&E) staining

Following UBD-VMR, bladders from C57Bl/6J female mice were harvested and placed in 4% paraformaldehyde (PFA) solution for 24 h, then 20% sucrose for cryoprotection. Following 24 h in sucrose, bladders were cut in half longitudinally and submerged in a mold with optimal cutting temperature compound (OCT Fisher Scientific Cat # 23-730-571), and stored at −80°C. Bladders were sectioned at 30 μm and mounted onto Poly-L-Lysine coated slides. After mounting, slides were baked at 40°C for 45 min and stored at 4°C. For hematoxylin and eosin (H&E) staining, bladder sections were first stained with filtered 0.1% Mayer’s Hematoxylin Solution (Sigma-Aldrich #MHS16) for 4 min followed by rinsing in cool running tap water (5 min) to stain for nuclei. Sections were placed in acid alcohol (10% acetic acid and 95% ethanol) for 1 min, then rinsed under cool running tap water for 1 min. Staining of the cytosol was done by staining the section in Eosin solution (Millipore Cat# HT110116)) for 1 min, followed by immediately dipping the slides in distilled water. Sections were dipped in increasing concentrations of ethanol (50% - 3 min, 70% - 3 min, 95% −30 s, and 100% - 1 min) before a final dip in xylene for 4 min. Cytoseal™ 60 (Thermo Scientific Cat# 23-244257) was applied to the section, followed by cover glass slips, and edges were sealed with clear nail polish. 4x and 10x magnification color images were captured with an Olympus IX73 inverted microscope and an iPhone X with an i-NTER LENS microscope adaptor (MR-6i) using the i-NTER SHOT2 app.

### Bladder histology quantification

A micrometer slide was used to set the scale of the microscope images. Quantitative image analysis was performed blinded to experimental treatment (DPI after CYP or saline) using ImageJ software and MS Excel to assess the thickness of the three layers of the bladder (urothelium, muscularis, and submucosa area) as previously described^29^. Four different measurements were taken of the urothelium and muscularis layers, and then averaged per section. For the submucosa area, the Image J freehand tool was used to measure the submucosa area and the total area of the entire bladder section, then input as a ratio. Three sections were quantified and averaged per animal for all layers.

### Direct injections of aCSF or CGRP8-37

Prior to behavior experiments, wildtype male and female C57BL/6J animals underwent stereotaxic cannula surgery, in which a stainless steel cannula (8.01mm long, 0.2 mm diameter; Microgroup-TE Connectivity) was lowered into the CeA (AP −1.45 mm, ML +/- 3.00 mm, DV - 4.20 mm) and fixed in place using dental cement as described previously^30^. Animals were allowed to recover for a week, prior to the start of testing. On test day, animals were under anesthesia (Isoflurane, 2.0%), while receiving a direct injection (1 μl) of either aCSF or CGRP_8-37_ via syringe pump (Harvard Apparatus, Pump 11 ver. 6.2). The injection was given over the course of 5 min at a rate of .200 µl/min The experimenter was blinded to treatment. Treatment was administered 40 min prior to abdominal mechanical sensitivity testing and 30 min prior to void spot testing.

### Behavioral Analysis

#### Abdominal mechanical sensitivity

The sample size for abdominal mechanical sensitivity was determined by an a priori analysis (G*Power) based on a previous pilot study. Abdominal mechanical sensitivity was conducted before (0 DPI) and after CYP treatment (1-20 DPI). At the start of behavioral testing, animals were placed in covered plexiglass enclosures (10mm L x 10 mm W x 16.5 mm H) on top of a wire mesh platform to habituate to the dimly lit room with white noise (60 dB). Using the up-down method^19,31^, a range of calibrated von Frey filaments (TouchTest; 0.02g (0.19mN), 0.04g (0.39mN), 0.08g (0.78mN), 0.16g (1.5mN), 0.32g (3.1mN), 0.64g (6.3mN), 1.28g (12.6mN), 2.56g (25.1mN)) were applied to the abdomen (∼0.5 cm away from the urethra) to determine the force required to elicit a withdrawal response. Starting from the 0.32g filament, different gauges of von Frey filaments were applied independently to the left and right regions of the abdomen. If there was not a withdrawal response, the filament gauge moved up by one (0.64g). If there was a withdrawal response, the filament gauge used moved down by one (0.16g). After a switch from withdrawal to no withdrawal response (or vice-versa), the test was repeated four more times.

Both the left and right abdominal regions were tested, then averaged together to calculate the final 50% withdrawal threshold.

### Void Spot Assay

A single mouse was placed in an enclosure (17.37cm L x 28.57cm W) lined with absorbent filter paper, allowing the mouse to move freely for 60 min. After the 60 min trial, the filter paper was collected and dried at room temperature for >12 hours. A UV blacklight flashlight (Quantum) was used to outline each urine void. The number of voids on each filter paper was counted and differentiated as either a “void” or a “microvoid.” Anything less than 0.37 microliter was considered a microvoid. To measure the surface area of each void, each filter sheet (with the voids outlined) had a 10 cm ruler taped to it (to set the scale) and was scanned using a high-resolution camera (iPad X). On a separate blank sheet of clean filter paper, known urine volumes were applied to calibrate quantification. ImageJ was used to calibrate pixels into cm using the markings on the 10 cm ruler placed on the void sheet. The surface area, mean, minimum, and max were measured in pixels/cm. The surface area for each void was summed and divided by the number of voids. Surface area to microliters was determined from the known urine volume control sheet. The surface area per void was converted to microliters per void with every 1 microliter being 2.7617 cm^2^. All void analysis and quantification was completed blinded to treatment condition. The number of fecal boli from each animal was recorded at the end of the voiding session.

### Urinalysis – Epithelial Cell Shedding

Urine (∼30 µl) was collected from either CYP or saline-treated female and male wildtype mice treated 0, 2, 7, 14, 21 DPI. Urine samples were immediately placed in −20 °C, then later stored at −80 °C until the pap staining of the urine was performed. Prior to embedding the urine samples onto microscope slides (Shandon Cytospin 3, GMI), 10 ul of urine was mixed with 40 ul of 1X PBS, transferred into a sample chamber, and spun at 1000 rpm for 6 min. Embedded samples on slides were fixed using acetic/alcohol fixative and dehydrated in 95% alcohol, followed by rehydration in water. Hematoxylin was incubated for 10 minutes and developed in running tap water. Sections were dehydrated in 95% alcohol and stained with brand OG-6 solution (22-050-211, Thermo Fisher Scientific, USA) for 2 min and washed with 95% alcohol. Next, EA-65 stain (22-050-211, Thermo Scientific, USA) was used for 10 min, followed by a dehydration step.

Clearing was done in xylene and a xylene-based mounting media was used to mount the slides, followed by imaging under a Panoramic Midi microscope (3DHISTECH Ltd, Hungary). All urinalysis and quantification were completed blinded to treatment condition.

### Ex vivo electrophysiology

#### Acute slice preparation

After 3 doses (across 5 days) of CYP treatment or saline control, at 6/7 DPI, *Calcrl*^Cre/wt^ x Ai14(RCL-tdT)-D^+/-^ male and female mice were decapitated and brains were rapidly extracted, placed in ice-cold cutting solution, and cut in coronal slices (250–300 μm) using a Leica VT1200 S vibrating blade microtome (Leica Microsystems Inc.). The cutting solution was composed of the following: 110 mM choline chloride, 25 mM NaHCO_3_, 1.25 mM NaH_2_PO_4_, 2.5 mM KCl, 0.5 mM CaCl_2_, 7.2 mM MgCl_2_, 25 mM D-glucose, 12.7 mM L-ascorbic acid, and 3.1 mM pyruvic acid, oxygenated with 95%/5% O_2_/CO_2_. The slices containing CeA were incubated at 25°C for at least 60 min in a holding chamber containing artificial CSF (ACSF) composed of the following: 125 mM NaCl, 2.5 mM KCl, 1.25 mM NaH_2_PO_4_, 25 mM NaHCO_3_, 2 mM CaCl_2_, 1 mM MgCl_2_, and 25 mM D-glucose. The slices were then moved to the microscope bath and recovered for at least 10 min at 33°C before recording. During incubation and recovery, the chambers were continuously oxygenated with 95%/5% O_2_/CO_2_.

### Whole-cell patch-clamp recordings

The recording chamber was perfused continuously with ACSF oxygenated with 95%/5% O_2_/CO_2_ (1 ml/min) and all recordings were performed at 33 ± 1°C. A recording chamber heater and an in-line solution heater (Warner Instruments) were used to control and monitor the bath temperature throughout the experiment. Recording pipettes (3.5-to 6.6-MΩ resistance) were filled with internal solution composed of the following: 120 mM potassium methyl sulfate, 20 mM KCl, 10 mM HEPES, 0.2 mM EGTA, 8 mM NaCl_2_, 4 mM Mg-ATP, 0.3 mM Tris-GTP, and 14 mM phosphocreatine with pH 7.3 using 5 M KOH and an osmolarity of ∼300 mosmol^−1^. Whole-cell current-clamp recordings were obtained from tdTomato-expressing CeA neurons located in the capsular division (CeC) or lateral division (CeL) in the right or left hemisphere. Cells were visually identified using an upright microscope (Leica DM6 F6) equipped with differential interference contrast optics with infrared illumination and epifluorescence. Recording electrodes were visually positioned in the CeC, guided by the distinctive fiber bundles and anatomic landmarks delineating its structure. Recordings were controlled using the Multiclamp 700B patch-clamp amplifier interfaced with a Digidata 1440A acquisition system and pCLAMP 10.7 software (Molecular Devices) on a Dell computer. Before forming a membrane-pipette seal, pipette tip potentials were zeroed. Whole-cell capacitance was derived from membrane time-constant calculation obtained from capacitance curve traces in current-clamp configuration. Spontaneously active cells were recorded gap-free in current-clamp configuration. Brief (5 ms) and prolonged (500 ms) depolarizing current of various amplitudes were injected from resting membrane potential to cells that were silent at rest, to elicit single and repetitive action potential firing, respectively. Liquid junction potentials were not corrected during recordings. All recordings were acquired at 20 kHz and filtered at 4 kHz. Recording sites were constructed using the mouse brain atlas as a guide (Paxinos et al., 2001). Position of cells ranged from −1.22 to - 1.70 mm of Bregma. Recordings were completed blinded to treatment condition.

### Electrophysiology Data Analysis

The sample sizes used in each experiment were based on previous studies of the CeA in nociception^32,33^. Cells were allocated into experimental groups based on saline-treated and CYP-treated groups, which were further categorized into left and right CeA. Electrophysiological data were analyzed using ClampFit 11.4 (Molecular Devices), Microsoft Excel, Mini Analysis (v. 6.0.8, Synaptosoft), and Prism (version 10.4.1, GraphPad Software Inc.). Single action potential properties were measured from the action potentials generated in response to a 5-ms depolarizing current injection. Current threshold for action potential generation was defined as the minimum current injection required to elicit an action potential. Action potential duration (APD) was measured at 100% repolarization to threshold potential. Rise time was defined as the time required for the membrane potential to reach peak voltage from threshold potential, and decay was defined as the time required for the membrane potential to repolarize from 90% of its peak to threshold potential. Voltage threshold, rise, decay and APD were manually measured from the traces. Fast afterhyperpolarization (fAHP) was measured as the peak of the repolarization’s downstroke. Medium afterhyperpolarization (mAHP) was measured as the slow peak occurring around 50 ± 25 ms from the action potential peak. Action potential peak voltage was measured as the most depolarized potential reached during a spike. For spontaneously firing cells, voltage threshold, rise, decay, APD, fAHP and mAHP were calculated using the methods described above. Rheobase was defined as the minimum current required to induce an action potential in response to a 500-ms depolarizing current injection for both late-firing and regular-spiking neurons. Latency to fire was calculated with 2× rheobase current injection and was defined as the time between current injection onset to action potential threshold. Voltage sag was calculated from the difference between the steady state and peak voltage responses to a 500-ms 500-pA hyperpolarizing current injection. Latency to first spike was used to classify cells as late-firing or regular-spiking neurons. Cells with latencies shorter than 100 ms (at baseline) or 90 ms (pain conditions) were classified as regular-spiking. Conversely, cells with latencies higher than 100 ms (at baseline) or 90 ms (bladder-pain condition) were classified as late-firing.

Accommodation of inter spike interval (ISI) was calculated from the ratio of the measurements obtained from the last and first action potential in response to a 500-ms depolarizing current injection at 2× rheobase. ISI accommodating cells were defined as cells with a ratio ≥1.5, whereas ISI non-accommodating cells had a ratio of <1.5. Input resistance (R_in_) was calculated using the average change in membrane potential in response to a ±20pA current injection of 500-ms duration. Whole-cell membrane capacitance was calculated by curve fitting the capacitive transients elicited by 10 sweeps of −80 to +100 pA current steps. The voltage transient’s exponential phase was used to derive the membrane time constant by a fitting method utilized by Levenberg-Marquardt algorithm in Clampfit software. Whole-cell capacitance is then calculated from the following equation:

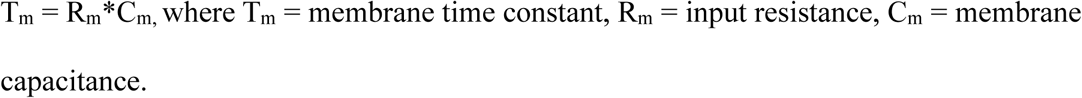

### In vivo Calcium Imaging

#### GCaMP6m virus and gradient refractive index (GRIN) lens implantation surgery

*Calcrl*^cre/wt^ male and female mice were anesthetized with isoflurane (3-5% for induction, 1.5-2% for maintenance) in the stereotaxic frame (model 1900, Kopf) for viral injection and GRIN lens implantation surgery. Mice were injected with 1000nL of diluted AAV.Syn.Flex.GCaMP6m.WPRE.SV40 (AAV9) virus (Addgene 100838; titer: 2.7×10^13^vg/ml, diluted to 6.75 x 10^12^ at a rate of 3.2 nL/sec using a 2.5 µl microsyringe (Hamilton, 7632) and a microsyringe pump system (UMP3-Micro4, WPI). Following injection, the needle was raised 200 µm for an additional 5 min to allow the virus to diffuse at the injection site, and then slowly withdrawn. Next, the GRIN lens (ProView™ 0.6mm x 7.3mm Integrated Lens,Inscopix Cat#1050-004413) was slowly lowered (200 µm/min until DV −2.00 mm; 100 µm/min until DV - 3.5 mm; 50 µm/min until DV −4.0 to −4.2) into the right (AP −1.42mm; DV −4.2mm; ML 3mm) or left CeA (AP −1.42mm; DV −4.2mm; ML −3mm). Metabond (Frontier Dental Supplies Cat# 945-0435AA) and bone screws (Stoelting Cat#51457) were used to secure the lens/baseplate in place. After surgery, the animal received 200 µl of 0.9% saline subcutaneously and 10 µl of 0.3 mg/ml buprenorphine (Covetrus Cat# 059122) intraperitoneally.

#### Measuring spontaneous and stimulus-evoked neural activity

Three to four weeks after implantation, a dummy miniscope (Inscopix Cat# 1050-003767) was attached to the baseplate of the implanted GRIN lens to habituate the animal to the miniscope and the covered plexiglass enclosure (10mm L x 10 mm W x 16.5 mm H) on top of a wire mesh platform for 30 min 1-2 days prior to imaging. After habituation experiments, the main protocol consisted of recording on 0 DPI (before CYP treatment), 2 DPI, 7 DPI, 14 DPI, and 21 DPI. At the start of each imaging session animals were under 3% isoflurane for ∼30 sec to mount the miniature microscope to the baseplate of the implanted GRIN lens. The microscope was tethered to a commutator system (Inscopix #1000-005088), which was connected to the Inscopix DAQ box (Inscopix nvoke Cat#: 100-004245). Prior to imaging, the field of view (FOV), gain, and LED light intensity were modified to observe optimal GCaMP6m fluorescence, and animals were habituated to the enclosure for 30 min. At the start of imaging, spontaneous neural activity was recorded for 5 min. Following spontaneous activity, a 6 min test trial was performed to assess Ca^2+^ transient activity before, during, and after, both innocuous and noxious von Frey filaments were applied to the abdomen of the mouse. At the start of the test trial, Ca^2+^ activity was recorded for 1 min (before innocuous test). Then, an “innocuous” von Frey filament (0.04g) was applied to the right and left abdomen in an alternating fashion for 1 min (during innocuous test), followed by 1 min of recording after the innocuous stimulus. The 0.04g filament was chosen as the “innocuous” filament because it almost never causes a noticeable behavioral response in naïve uninjured mice. Next, Ca^2+^ activity was recorded for 1 min (before noxious stimulation). Then, a “noxious” von Frey filament (2.56g) was applied to the right and left abdomen in an alternating fashion for 1 min (during noxious), followed by 1 min of recording after the noxious stimulus (after noxious). The 2.56g filament was chosen as the “noxious” filament because it is a filament that causes a response in almost all naïve, uninjured mice.

#### In vivo Imaging Data Analysis

All miniature fluorescent microscope videos were recorded at a frame rate of 10 or 20 Hz, using between 1.4 and 2.0 LED intensity. All Ca^2+^ imaging movies were pre-processed using the Inscopix Data Processing Software (IDPS). Using IDPS, movies were down-sampled, motion corrected, and bandpass-filtered. Following pre-processing, individual neurons and their activity traces were extracted using the PCA-ICA algorithm. After PCA-ICA, neurons were further verified manually by pixel size and signal-to-noise (SNR) ratio because in our use of this system fully automated PCA-ICA analysis sometimes identified areas that appeared to be noise rather than true cellular GCamP6m signal.

### General Statistics

All data analyses were conducted blinded to treatment. UBD-VMR data: Two-way (mixed effects) ANOVA followed by Tukey’s post hoc test were used to analyze data. Bladder histology quantified data: One-way ANOVA followed by Tukey’s post hoc test were used to analyze data. Pharmacological data: Two-way RM or mixed effects ANOVA followed by Šídák’s multiple comparisons test were used to analyze data. Electrophysiology data: Unpaired t-test, ANOVA followed by Tukey’s post hoc test were used to analyze data. *In vivo* calcium imaging data: Two-way (mixed effects) ANOVA followed by Šídák’s multiple comparisons post hoc test were used to analyze data. Statistical significance was determined at the level of *p* < .05. Asterisks denoting *p* values include ∗*p* < .05, ∗∗*p* < .01, ∗∗∗*p* < .001, and ∗∗∗∗*p* < .0001. All data are presented as the mean ± SEM.

## Results

### CYP induces hypersensitivity in UBD-VMR 7 and 14 DPI, without disrupting bladder histology

To study the long-term effects of CYP-induced injury on bladder pain physiology, we measured the UBD-VMR in female mice that were treated with either saline (0 DPI) or CYP at 2, 7, 4, and 21 days post-injury (DPI) timepoints (**Fig. 1A**). CYP induced statistically significant bladder pain-like physiological responses in the UBD-VMR test at 7 and 14 days post-injury (**Fig. 1B**). Mixed-effects analysis revealed a significant effect of time (DPI) (p= 0.0005) and pressure (p= 0.0077). Tukey’s multiple comparisons test reveals a significant increase in normalized VMRs in both 7 and 14 DPI groups when compared to the saline-treated group (0 DPI) for both 30 mmHg and 60 mmHg pressures. Next, we measured potential bladder histology changes after prolonged treatment with CYP, when compared to saline-treated animals (**Fig. 1C-D**). To measure histological changes, bladders underwent H&E staining, and the thickness of the urothelium and muscularis layers of the bladder were measured from saline-treated mice (0 DPI) and mice that received CYP 2-21 DPI. We also measured the submucosa area relative to the total area of the bladder. Our qualitative observation was that some bladder tissue appeared more compact than others. Once unblinded, we observed the more compact tissue was observed more from tissue from saline-treated mice than in bladders from mice that received cyclophosphamide (**Fig. 1C**). However, once quantified, we observed similar bladder histology across all groups (**Fig. 1D**).

The mean urothelium thickness for the saline group (0 DPI) was 37.5 μm ± 5.24 SEM. One-way ANOVA revealed there was no significant difference in muscularis thickness among the different groups (p= 0.138). The mean submucosa ratio for the 0 DPI group was 32.0% ± 5.09 SEM. In the CYP-treated animals, the submucosa ratio was 33.4% ± 3.46 SEM in the 2 DPI group, 35.9% ± 3.58 SEM in the 7 DPI group, 37% ± 2.15 SEM in the 14 DPI group, and 34.2% ± 3.20 SEM in the 21 DPI group. One-way ANOVA revealed there was no significant difference in submucosa ratio among the different groups (p= 0.8863). In a separate group of animals, we measured the amount of epithelial cell shedding in urine from CYP and saline-treated mice (**Supplemental Fig. 1**). We found an overall increase in epithelial cell shedding in female mice compared to males. In males, we observed increased epithelial cell shedding at 14 DPI in CYP-treated mice, when compared to saline controls.

### CGRP-R antagonist CGRP_8-37_ attenuates the development of abdominal mechanical hypersensitivity when administered in the right CeA

To determine the extent to which CGRP receptors are important in the development of persistent bladder pain, we pharmacologically inhibited CGRP-R activity in the right or left CeA prior to observing bladder pain-like behavior in male and female mice. For this study, mice received direct injections of either artificial cerebrospinal fluid (aCSF) or the peptide antagonist CGRP_8-37_ prior to each abdominal von Frey test and each void spot assay test 0-21 days post injury (**Fig. 2A**). In the left CeA, CGRP_8-37_ had no significant effect on abdominal mechanical sensitivity when compared to mice that received aCSF (**Fig. 2B**). In both aCSF and CGRP_8-37-_ treated animals, mice displayed significant abdominal mechanical hypersensitivity by 6 DPI, when compared to 0 DPI (i.e., before CYP treatment) consistent with our observations in the UBD-VMR assay (**Fig. 1B**). Two-way ANOVA revealed a significant effect of time (p< 0.0001), but not CeA treatment (p= 0.3216). Furthermore, Šídák’s multiple comparisons test revealed no significant differences between treatment groups at any of the different timepoints (i.e., 0-20 DPI). However, CGRP_8-37_ in the right CeA significantly attenuated the development of persistent bladder pain (**Fig. 2C**). Two-way ANOVA revealed a significant effect of time (p< 0.0001) and treatment (p= 0.0089). Šídák’s multiple comparisons test revealed CGRP_8-37_, when injected into the right CeA, significantly increased the 50% withdrawal threshold at 6 DPI (p= 0.0280) and 21 DPI (p= 0.0465), when compared to aCSF treatment. These data suggest that inhibition of CGRP-R activity in the right CeA alone can attenuate the development of persistent bladder pain-like mechanical hypersensitivity.

**Figure 2:**
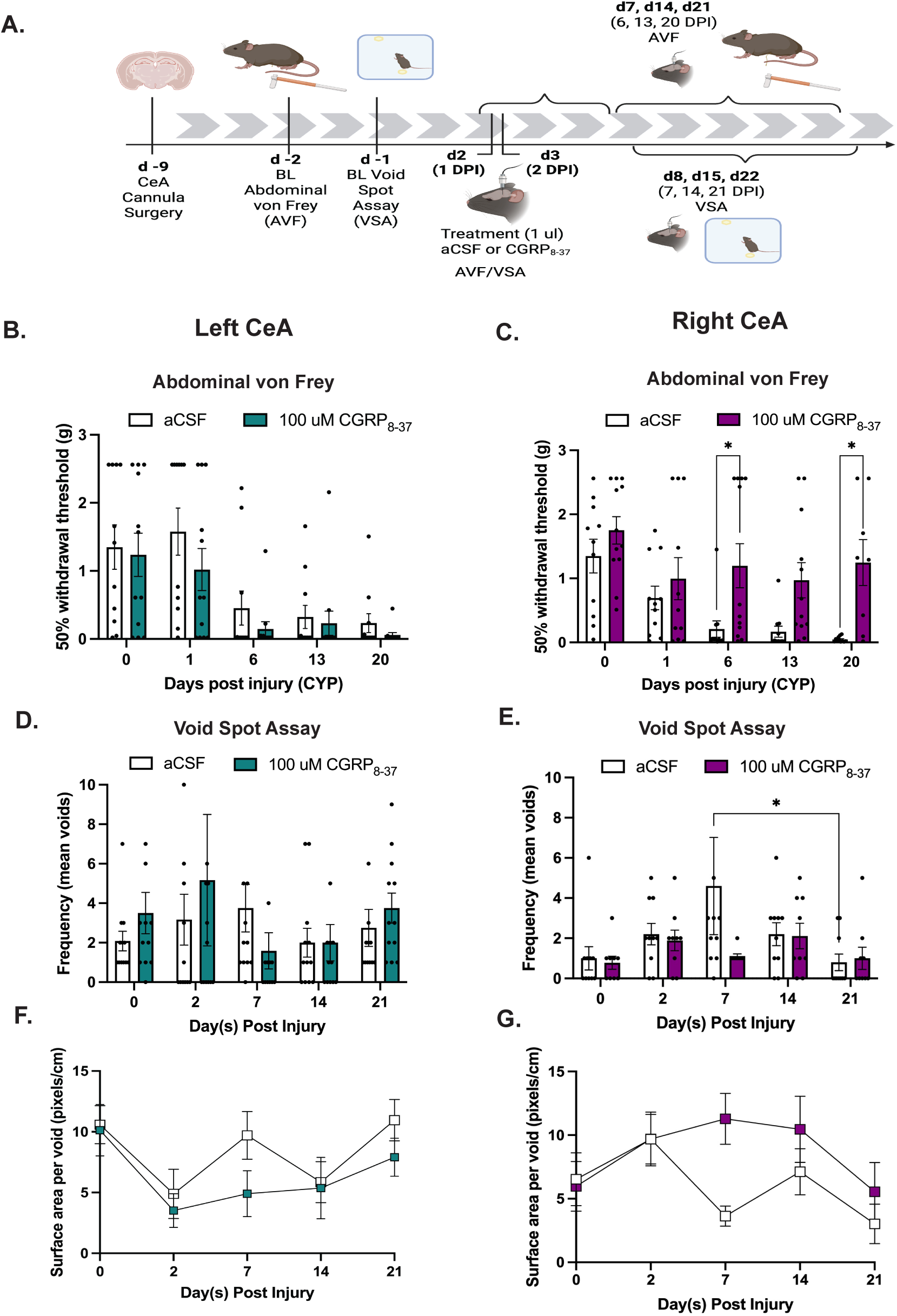
CGRP_8-37_ attenuates the development of persistent bladder pain in the right CeA only. **A)** Schematic of experimental timeline. **B-C)** Abdominal mechanical sensitivity of mice treated with either CGRP**_8-37_** or aCSF in the **B)** left CeA, n=12/group or **C)** right CeA, n=8-12/group. Results are presented as mean ± SEM. Asterisks represent results from Šídák’s post hoc test. Šídák’s multiple comparison test revealed a significant difference between treatment groups (*, p<0.05). **D-E)** Average frequency of voids in mice treated with either CGRP**_8-37_** or aCSF in the **D)** left CeA, n= 12/group or **E)** right CeA, n=10-12/group. Asterisk represents results from Šídák’s post hoc test (*, p< 0.05). **D-E)** Results are presented as the frequency of voids from each mouse during the void spot assay. **F-G)** Surface area per void during the void spot assay from mice treated with either CGRP**_8-37_** or aCSF in the **F)** left CeA, n=12/group or **G)** right CeA, n= 8-12. Asterisks represent results from Šídák’s post hoc tests comparing treatment groups (*, p< 0.05).

### Changes in voiding behavior are observed 7 days post injury but these changes are not impacted by CeA CGRP-R manipulation

Patients with IC/BPS often report frequent urination^34^. Therefore, in the same group of mice, we studied the effect of CGRP_8-37_ on voiding behavior 0-21 DPI (**Fig. 2D-G**) and fecal boli production (see **Supplementary** Fig. 2). CYP did not have a significant effect on the frequency of voids in aCSF-treated animals (Supplementary Fig. 2E). CGRP_8-37_ administered in the left CeA had no effect on the frequency of voids when compared to mice that received aCSF (**Fig. 2D**). Two-way ANOVA revealed no significant effects of time (p= 0.5065) or treatment (p= 0.6297). Šídák’s multiple comparisons test revealed no significant differences in the frequency of voids between treatment groups 0-21 DPI. Similarly, in the right CeA, CGRP_8-37_ had no significant effect on the frequency of voids when compared to mice that received aCSF (**Fig. 2E**). Two-way (mixed-effects) ANOVA revealed no significant effects of time (p = 0.2113) or treatment (p= 0.1911). However, Šídák’s multiple comparisons test revealed a significant increase in voiding frequency at 7 DPI, compared to 21 DPI, in mice that received aCSF in the right CeA (p= 0.0454).

Accompanying the increase in frequency of voiding by patients with IC/BPS is a reduction in urine voided per event^35^. Essentially, the pain associated with filling leads to an increase in urination with each bought having less volume than a regular void. Therefore, we also measured the surface area of each void spot. In the left CeA, CGRP_8-37_ had no significant effect on surface area per void, when compared to aCSF (**Fig. 2F**). Two-way ANOVA revealed a significant effect of time (p = 0.0071) but no effect of treatment (p= 0.1493) on surface area per void. Although not significant, at 7 DPI aCSF in the left CeA displayed an increase in void surface area when compared to CGRP_8-37_. In the right CeA, Two-way ANOVA revealed no significant effects of time (p= 0.0650) or treatment (p= 0.0608) (**Fig. 2G**). These data suggest a minimal change in voiding behavior at 7 DPI that can be attenuated by blocking CGRP-Rs.

### CYP increases the firing rate of CGRPR+ neurons in the right CeA

To assess bladder-pain-induced neuronal excitability changes between the left and right CeA’s CGRP-R positive populations, we recorded neuronal excitability from acute brain slices 6 or 7 DPI. We performed whole-cell current-clamp recordings on visually identified CeC CGRP-R^+^ cells by Cre-dependent expression of tDTomato (**Fig. 3A**). A total of 61 cells were recorded from 21 mice (9 mice from the CYP group and 12 mice from the saline-treated control group) (**Fig. 3B**). All the recorded cells were from capsular (CeC) and lateral (CeL) sub-regions of the CeA, and a majority were from posterior (bregma −1.70 + 0.3 mm) CeA slices (**Fig. 3L**). Based on their latency to fire, the CeA neurons are known to exhibit heterogeneous firing identities, such as the late-firing (LF), regular-spiking (RS), and spontaneously firing (spon) types^36,37^. To evaluate whether CGRP-R^+^ neurons exhibited such heterogeneity, we injected a depolarizing step current at twice the rheobase for 500 ms. As expected, we observed that CGRP-R^+^ neurons exhibited late-firing (LF), regular-spiking (RS), and spontaneously firing (spon) phenotypes (**Fig. 3C**). LF and RS types remained silent at resting potentials, while the spon type fired at resting potentials without any stimulating current injection. Of the total number of cells recorded, the majority of the neurons were regular-spikers (50/61; 81.9%), followed by a minority late-firing type (9/61; 14.7%), and rarely occurring spontaneous type (2/61; 3.2%). We wanted to evaluate whether there was a difference in the distribution of CGRP-R^+^ cellular firing identities between the hemispheres of saline-treated mice and CYP-treated mice. Population distribution analysis showed that the distribution of firing phenotypes did not differ between the left and right CeA in both saline and CYP groups (p > 0.7318) (**Fig. 3D**).

**Figure 3.**
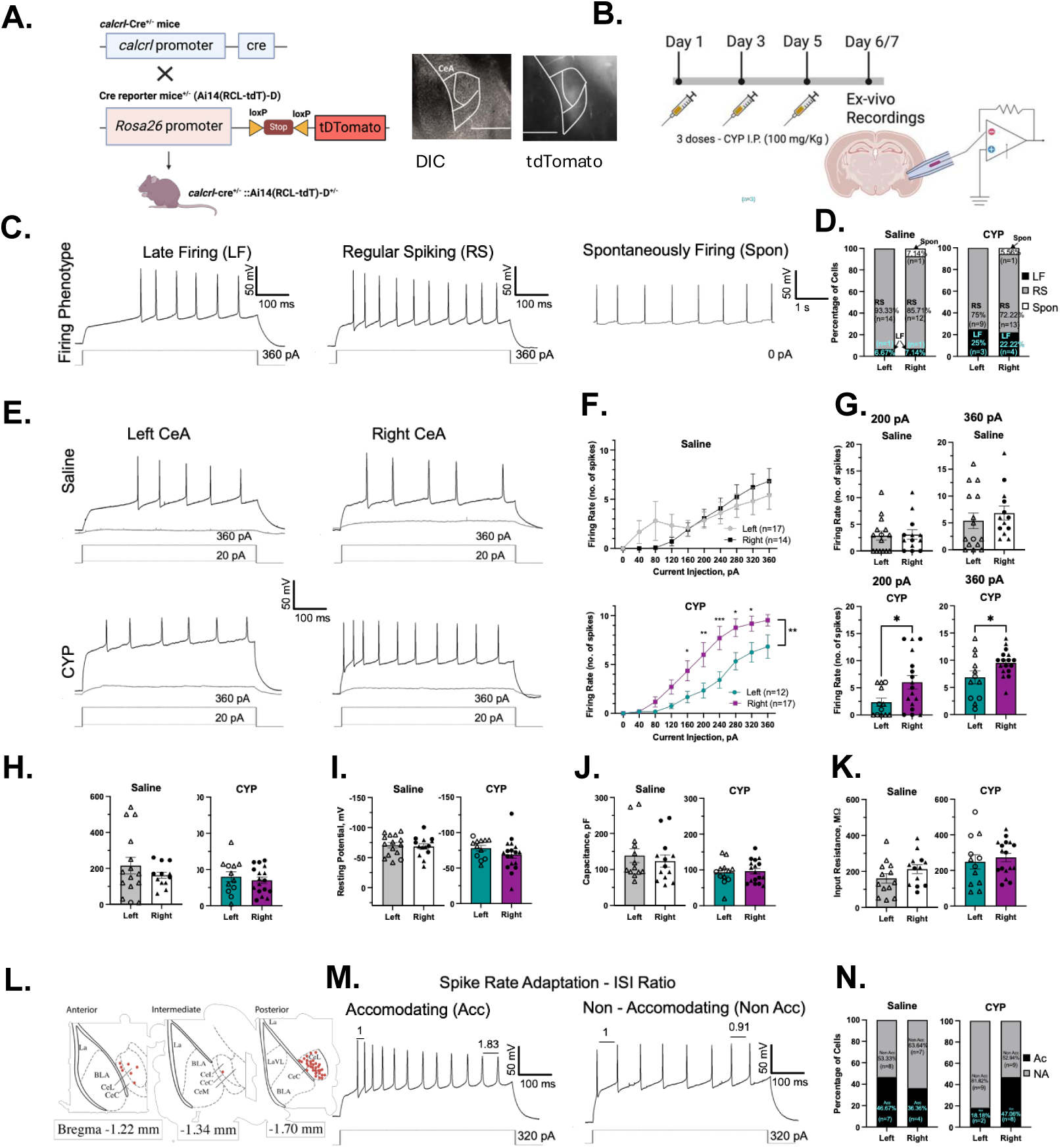
The right CeA CGRPR^+^ neurons show increased firing after CYP-induced bladder pain. **A.)** Schematic of CGRPR^+^ neuronal targeting strategy (top left). Representative image of CeA containing brain slice under DIC microscopy (left image, scale bar - 600), and epifluorescence (right image) showing tDTomato expression indicative of CGRPR^+^ neurons. **B)** Schematic of the timeline of CYP dosage and *ex-vivo* electrophysiology. **C**) Representative traces of late-firing (LF), regular-spiking (RS), and spontaneously firing (spon) phenotypes. **D)** Proportion of firing phenotypes. **E)** Representative traces repetitive firing from left and right CeA in saline group (top panel) and CYP group (bottom panel). **F)** Input-output curves plotted as number of spikes between left and right CeA in saline group (top) and CYP group (bottom). **G)** Number of spikes elicited by 200pA and 360pA between left and right CeA in saline group (top) and CYP group (bottom). **H)** Resting membrane potential between left and right CeA in saline (left) and CYP group (right). **I)** Rheobase between left and right CeA in saline (left) and CYP group (right). **J)** Whole-cell capacitance between left and right CeA in saline (left) and CYP group (right). **K)** Whole-cell input resistance between left and right CeA in saline (left) and CYP group (right). **L)** Anatomical location of recorded neurons across the anterior-posterior axis (rostro-caudal axis) of the CeA. **M)** Representative traces of accommodating and non-accommodating types of spike rate adaptation phenotypes. **N)** Proportion of spike rate adapting phenotypes. All data are presented as mean ± SEM, and error bars represent SEM.**p<0.01, *p<0.05. Individual data points belonging to males and females are denoted with triangle and circle symbols, respectively.

Next, to evaluate the neuronal input-output function in normal and CYP groups, we injected depolarizing step currents of increasing amplitudes into the neurons. In the CYP group, depolarizing amplitudes produced a statistically significant increase in the number of action potentials only in the right CeA (p = 0.0015) but not left CeA (**Fig. 3E-G**). However, no differences were observed in rheobase and passive membrane properties (i.e., resting potential, capacitance, and input resistance) between the hemispheres across CYP and saline groups (**Fig. 3H-K**). Furthermore, no changes were observed in single action potential properties such as the spike amplitude, action potential duration, rise time, decay time, fAHP peak, mAHP peak, current threshold, voltage threshold, or in hyperpolarization-activated sag potential (**Supplemental Fig. 3 A-M**).

To evaluate whether the increase in the firing rate in the right CeA in the CYP group is due to any difference in spike-frequency adaptation properties between the hemispheres, we measured the inter-spike intervals (ISI) in current-injected repetitive spikes. Based on the ISI ratio between the first and the last intervals, we observed accommodating and non-accommodating phenotypes (**Fig. 3M**). Neurons exhibiting an ISI ratio of >1.5 were considered accommodating, and others were considered non-accommodating. The distribution of the adaptation phenotypes between left and right sides was not significantly different across the groups (**Fig. 3M-N**) (p > 0.2264). Overall, these results show that CYP-induced bladder sensitization in mice increases the neuronal excitability of the CGRP-R^+^ neurons of the right CeA only in terms of the input-output function. This bladder-sensitization-induced change in the output function is not due to intrinsic passive neuronal properties, single action potential properties, or spike rate adaptation properties.

### Neural activity (Ca activity) in CGRP-R+ neurons is increased in the right CeA shortly after injury is induced

Lastly, we sought to determine if CGRP-R *in vivo* neural activity varied between hemispheres as bladder injury transitioned from the acute to persistent state. To achieve this, we performed *in vivo* single-photon calcium imaging to indirectly measure neural activity in awake, behaving animals. *Calcrl*^Cre/wt^ mice were injected with AAV9-Syn-FLEX-GCaMP6m into the right or left CeA (**Fig. 4A**). Shortly after injections, a GRIN lens was implanted into the CeA to allow for miniscope visualization of changes in calcium. Calcium imaging sessions were conducted before (0 DPI) and at different time points after CYP treatment (0-21 DPI) (**Fig. 4B**). Change of fluorescence was measured before, during, and after an innocuous and noxious mechanical stimulus was applied to the abdomen of the animal. In naïve animals (0 DPI), we found the number of significant calcium events to be similar before, during, and after the animals were presented with the innocuous stimulus (**Fig. 4D**). Mixed-effects analysis revealed there was no significant effect of time (p= 0.6465) or hemisphere (p= 0.0662). However, a significant effect was observed in the interaction of time x hemisphere (p< 0.0001). During and after the presentation of a noxious stimulus, Šídák’s multiple comparisons test revealed a significant increase of CGRP-R activity in the right CeA during (p= 0.0078) and after (p=0.0023) the stimulus, when compared to the left CeA. Shortly after CYP was induced (2 DPI), there was a significant increase of CGRP-R activity in the right CeA when compared to the left CeA. Mixed-effects analysis revealed a significant effect of time (p< 0.0001), hemisphere (p< 0.0001), and interaction (time x hemisphere) (p< 0.0001). Šídák’s multiple comparisons test revealed an increase in calcium events before, during, and after the application of both innocuous and noxious stimuli (**Fig. 4E, left**). At 7 DPI, CGRP-R activity was similar across hemispheres (**Fig. 4E, middle**). Mixed-effect analysis revealed no significant effects of time, hemisphere, or interaction. At 21 DPI, the number of events in the right CeA was increased after the presentation of both innocuous and noxious stimuli (**Fig. 4E, right**). Mixed-effects analysis revealed a significant effect of hemisphere (p=0.0047), but not time or interaction. Šídák’s comparisons test revealed a significant increase of CGRP-R activity in the right CeA after the presentation of a noxious stimuli (p=0.0059), when compared to the left CeA. In the left and right CeA, the majority of the observed CGRP-R positive cells had no significant change in activity at 0 and 2 DPI when a noxious stimulus was presented, compared to 1 min prior (**Fig. 4G**). However, there were still a population of CGRP-R positive cells in both the left and right CeA that were both excited and inhibited following the noxious stimuli applied to the abdomen.

**Figure 4.**
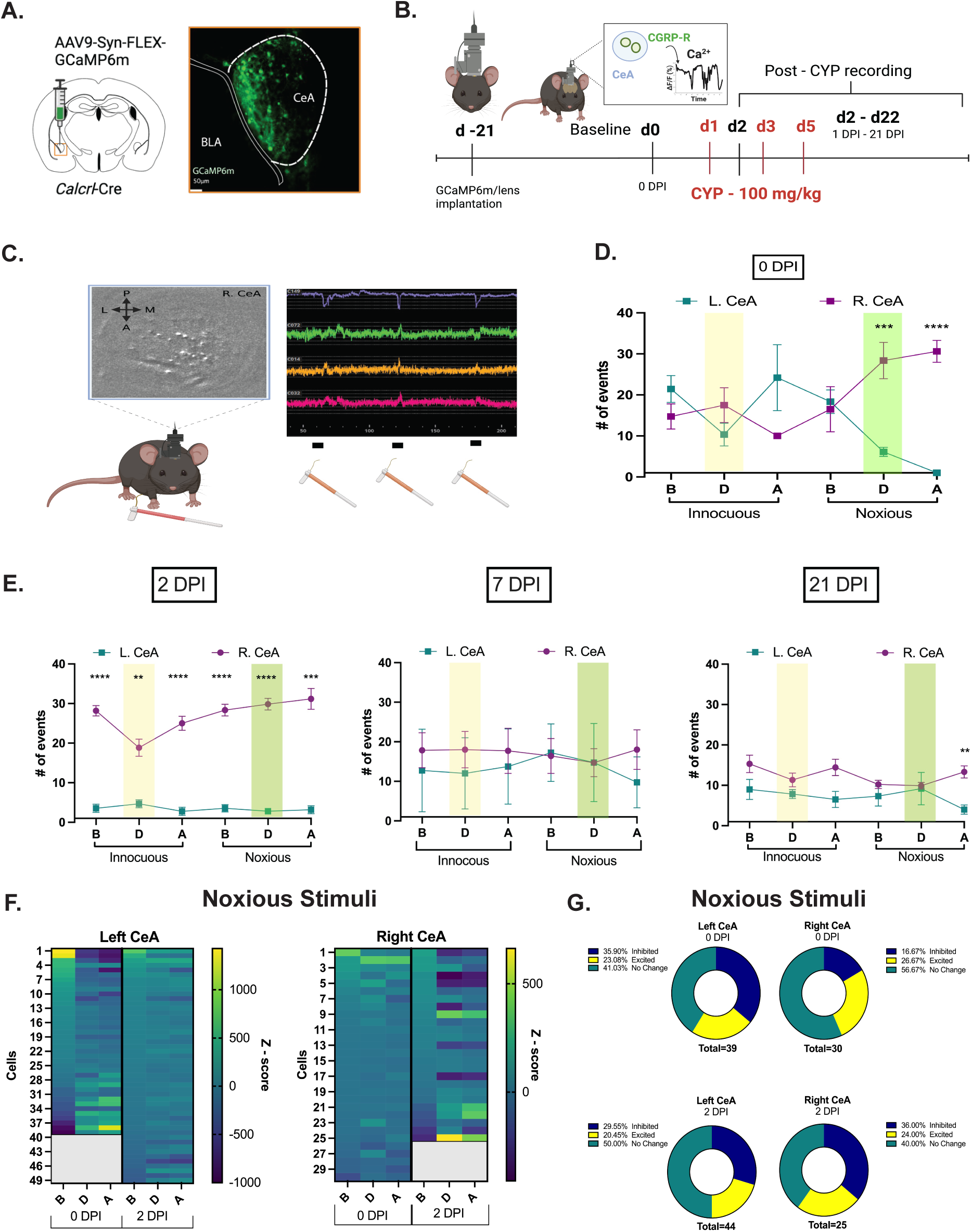
The right CeA CGRPR^+^ neurons show increased fluctuations in calcium events shortly after CYP is induced (2 DPI), when compared to the left CeA. **A-B)** Experimental design. **A)** Calcrl-cre mice were injected with GCaMP6m and implanted with a GRIN lens in the left or right CeA. Calcium imaging was conducted 0-21 DPI. **C)** Representative images and traces of stimulus-evoked activity. Black bars represent the time a von Frey filament was applied to the abdomen. **D)** Calcium responses in the left or right CeA in naïve (pain-free) animals. Y axis represents: # of calcium events (both negative and positive Ca fluctuations). B = before stimulus, D= during stimulus, A= after stimulus. Two-way ANOVA (mixed-effects model) shows a significant effect of time x hemisphere (p< 0.0001, ****). Šídák’s multiple comparisons test revealed significant hemispherical differences during (p= 0.0002, ***) and after (p<0.0001, ****) the noxious stimulus was applied. **E)** Calcium responses in CYP-treated animals, 0-21 DPI. Šídák’s multiple comparisons tests revealed significant hemispherical differences at 2 DPI (p< 0.0001, ****). **F)** Heatmaps representing the average Ca activity for cells before **(B)**, during **(D)**, and after **(A)**, the application of a noxious stimulus at 0 and 2 DPI. **G)** Represents the population of cells that were “unchanged”, excited, or inhibited upon the application of the noxious stimulus to the abdomen. 2-3 mice were used per group; graphs represent 25-44 cells.

## Discussion

The present study demonstrates that CYP is able to induce hypersensitivity up to 14 days post injury as displayed in the UBD-VMR assay and up to 21 days post injury when measuring abdominal mechanical sensitivity. These findings validate the use of CYP to observe the transition from acute to persistent bladder pain-like changes in mice. Furthermore, we observed this persistent hypersensitivity despite no significant changes in bladder histology. Prior to this study, our lab demonstrated that the neuropeptide CGRP contributed to CeA lateralization in the context of bladder pain but these studies were completed only up to 7 DPI^30^. In the CeA, CGRP is released from projection neurons whose cell bodies are in the parabrachial nucleus (PBN)^38^.

Here we demonstrate that CGRP-Rs also contribute to the modulation of persistent bladder pain in that inhibiting CGRP-R activity in the right CeA attenuates the development of persistent bladder pain. In contrast, pharmacological inhibition of CGRP-R activity in the left CeA led to persistent CYP-induced abdominal mechanical hypersensitivity that was no different from the aCSF-control treatment. Our electrophysiology findings reveal that once CYP damage occurs, firing frequency is increased in CGRP-R positive cells in the right CeA, when compared to the left CeA. Lastly, in our *in vivo* calcium imaging study, we demonstrate an increase in calcium transients in CGRP-R positive cells in the right CeA both before CYP and shortly after CYP treatment is initiated. These data further confirm the pronociceptive function of CGRP-R in bladder pain in the right amygdala. To our knowledge, this is the first study to assess the role of CGRP-R positive CeA neurons on the development of persistent bladder pain in mice.

This study sought to assess the long-term effects of CYP treatment in mice. CYP is a chemotherapeutic agent whose principal toxic metabolite, acrolein, is recognized for causing inflammation and damage to the bladder epithelium^39^. In humans, CYP treatment is associated with the induction of hemorrhagic cystitis, increased voiding frequency, and visceral nociceptive responses^24,40^. CYP has been used as a validated rodent model for overactive bladder and IC/BPS for years^25,41,42^. One large systemic dose of CYP is often used for acute models, while multiple systemic injections of lower doses have been used for chronic/persistent models of IC/BPS^43–45^. To our knowledge, the long-term effects of CYP on bladder pain-like behavior beyond 15 DPI have not been observed in mice prior to this study. Although similar effects to what we have observed have been seen previously at 7 DPI^46^. A study by DeBerry et al. found that while nociceptive responses to bladder distensions were observed at 7 DPI, signs of cystitis had resolved^46^. The long-term effects of CYP treatment have been observed in rats where histopathological changes in the bladder and changes in voiding behavior (e.g. urination interval and maximum voiding pressure) were observed 30 days and 45 days after the first day of CYP treatment^47^. Here, we observed an increase in the UBD-VMR at 7 DPI, which corroborates previous studies in our lab where CYP increased pain-like responses to UBD 24 h to 48 h following the final injection of CYP^18,19^. Next, we observed hypersensitivity up to 14 days post-injury during UBD-VMR. Although not significant, we observed a partial return to baseline at 21 DPI, showing that the effect on UBD-VMR is likely not permanent. We also showed consistent mechanical hypersensitivity from 6 DPI through 21 DPI. Despite the hypersensitivity observed during UBD-VMR and abdominal sensitivity testing, the same mice did not display noticeable histological changes in the bladder following CYP treatment. These data were surprising given that other studies have shown CYP to increase urothelium and submucosa thickness^48,25^. This could partially be due to different CYP dosing procedures or higher variability observed between subjects in our study. We did not stain bladders for principal proinflammatory markers so it is difficult to fully conclude that CYP does not induce any inflammation in the bladder. Similar to other studies^45,48–50^, we qualitatively observed that bladders from CYP-treated mice, particularly at 14 and 21 DPI, had pronounced edema in the submucosa and tissue separation when compared to bladders from the saline-treated mice.

Next, we tested the role of CGRP-Rs in the development of persistent bladder pain. CGRP has been found to modulate nociceptive signaling, promote inflammatory responses, and facilitate vasodilatory processes^51^. CGRP receptors are present in high densities within the CeA^52^. We demonstrated that when CGRP activity in the “antinociceptive” left CeA is pharmacologically inhibited, male and female mice still display severe abdominal mechanical hypersensitivity although the effects were not statistically different from aCSF-treated mice. The failure to see statistically significant effects in the left CeA may be due to the fact that these mice already showed hypersensitivity in the assay making it more difficult to show more pain-like behavior on top of the CYP hypersensitivity. In contrast, when CGRP activity in the “pronociceptive” right CeA is pharmacologically inhibited, abdominal mechanical hypersensitivity was attenuated. This corroborates recent studies in our laboratory that demonstrated that CGRP_8-37_ administered into the right CeA decreased pain-like responses at 7 DPI^19^. This observation of CGRP inhibition in the right CeA attenuating pain, and CGRP inhibition in the left CeA having mild pronociceptive effects, is consistent with some of the literature reporting amygdalar CGRP effects on pain-like behavior. In a rat arthritis pain model, CGRP_8-37_ administered into the right CeA inhibited mechanical hypersensitivity of the hindpaw and audible and ultrasonic vocalizations^7^. In naïve rats, CGRP_8-37_ administered into the left CeA inhibits the antinociceptive effects of CGRP on thermal and mechanical withdrawal thresholds^53^. In a rat model of neuropathic pain, CGRP_8-37_ administered into the right CeA decreased mechanical sensitivity in the von Frey and paw compression test^54^. In contrast to our data, a recent study from our lab found that CGRP8-37 in the left and right CeA increased cold sensitivity in the right SNI-treated hindpaw only, and had no significant effect of mechanical sensitivity in either SNI or chemotherapy-induced peripheral neuropathy (CIPN) model^55^. This suggests that CGRP functional lateralization is partially dependent on the pain model, the side of injury, and the behavioral endpoint.

Several studies have reported that CYP-induced cystitis increases voiding frequency and decreases urine volume/surface area per void^45,56–59^. However, in our hands, CYP did not have a robust effect on voiding behavior in that there were no significant differences observed in the aCSF CeA group at 0 DPI vs 2, 7, 14, or 21 DPI CYP, when both aCSF animals injected in the right and the left CeA were combined (**Supplemental Fig 2E**). Regarding the effects of CGRP_8-37_, no direct effect on voiding behavior was observed, as there were no significant differences between treatments. There was an increase in voiding frequency in mice treated with aCSF at 7 DPI when compared to 21 DPI, and CGRP_8-37_ into the right CeA increased the urine surface area per void. One explanation of less voiding frequency being observed at 21 DPI is a decreased sensitivity of CYP on the bladder, indicating the effects of CGRP signaling are diminishing by 21 DPI. This would also explain the decreased trend in pain-like responses to UBD that we observed at 21 DPI, when compared to 7 and 14 DPI. We selected a 1 hr void spot assay duration due to previous studies that suggested a short duration of action of CGRP_8-37_ with a peak at 45 min after treatment in the CeA^19^. It is possible that this 60 min window was not long enough to capture the effects of CYP on voiding. One study found that 2-4 hours was the most optimal time length for the assay^60^. The same study also suggested that time of day plays a major role in voiding. Our void spot assay was conducted over the span of 1 hr and in each test day and about 4 h elapsed between the first cohort of animals and the last cohort of animals tested in our study. It is possible that these factors contributed to our unexpected results.

Next, we measured how bladder-pain changes neuronal excitability of CGRP-R^+^ cells in the left and right CeA. For this particular study, we recorded neuronal activity using slice physiology and indirectly using single photon imaging. First, we measured excitability from acute brain slices from control animalsnd bladder-sensitized mice at 6/7 DPI.

We found CGRP-R firing to be similar in the left and right CeA of control animals. As hypothesized, in CYP animals, we observed an increase in firing frequency of CGRP-R^+^ cells in the right CeA, when compared to the left CeA. These data from control mice is consistent with our recent publication showing no hemisphere specific differential firing rates of left and right CeA neurons in response to exogenously applied CGRP^19^. No previous study has measured CGRPR^+^ neuronal excitability in any acute pain model. We show that in a bladder pain model there is increased excitability in CGRPR^+^ neurons in the acute phase of injury in the right CeA. A recent study, however, measured CGRPR+ neuronal excitability in a model of neuropathic pain, and found no change in firing rate at the chronic phase^33^. Nevertheless, earlier studies have shown arthritic pain associated increased neuronal excitability and synaptic plasticity in unmarked CeC neurons that were attenuated with CGRP1 receptor antagonists (CGRP^8–37^ and BIBN4096BS) in the right CeA^7^. Similarly, in a model of inflammatory pain, in CGRP knock out rats, pain associated synaptic plasticity was attenuated in unmarked CeC neurons, in the right side^61^. Both of these earlier studies were conducted at the acute phase of injury and in molecularly unidentified cell types. These earlier studies utilized pharmacological approaches and implicated CGRP-R’s role, indirectly, in potentiating central amygdala neuronal excitability and inducing synaptic plasticity. The advent of a CGRPR+ neuron-specific Cre line has proven to be an invaluable tool for targeting and studying this specific population in physiological investigations. Overall, we found that *ex-vivo* recordings from bladder sensitized mice suggest that the right CeA CGRPR+ neuronal sensitization can be detected at 6/7 DPI with exogenous current injection.

Second, we measure if CGRP-R neural activity varied between hemispheres as bladder pain transitioned from the acute to persistent state. We utilized *in vivo* calcium imaging, as an indirect measure of neural activity, when animals were mechanically stimulated in the abdomen. At 0 DPI, we observed an increase in CGRP-R activity once the noxious stimulus was applied. These data suggest that the contribution of CGRP-R signaling could be partially dependent on the severity of the stimulus. Surprisingly, we noticed an overall significant increase in CGRP-R activity in the right CeA shortly after CYP was induced (2 DPI), but no hemispherical differences were observed at 7 DPI. This is different from what was observed in a model of neuropathic pain, where there was an increase in general left CeA activity shortly after SNI, but an increase in right CeA by 14 DPI^62^. The main differences between their study and our study, is the model of pain, and the fact that they were observing overall CeA activity between the left and right, while we were specifically studying CGRP-R positive cells. Therefore, two possibilities exist: 1) the shift in CeA dominance is not observed in the transition from acute to persistent bladder pain and 2) CGRP/CGRP-Rs do not greatly contribute to the left to right CeA shift in dominance. Future studies should be conducted to assess the hemispherical differences in the transition from acute to persistent bladder pain in other CeA subtypes.

Seeing no hemispherical differences in calcium signals from CGRP-R positive cells at 7 DPI was surprising because of the difference seen between hemispheres at 7 DPI in our slice electrophysiology study. In awake *in-vivo* calcium recordings, bladder sensitization-induced activated PBN projection neurons to the CeA could synapse with local interneurons. These interneurons could then provide potentiated feedforward inhibitory modulations to CGRP-R^+^ neurons at 7 DPI. *Ex-vivo* recordings suggest that the altered intrinsic excitability of CGRP-R^+^ neurons after CYP-induced bladder pain persists even at 6/7 DPI. However, this increased excitability is only observed when depolarizing currents are exogenously injected into a single CGRPR^+^ neuron. In contrast, during in *in-vivo* stimulation the local network is engaged by an exogenous stimulus. Another possibility is that during *in vivo* stimulation, the excitatory synaptic transmission does not produce large enough excitatory post synaptic potentials at 7 DPI to evoke an action potential, which in turn is reflected as low calcium influx. Further electrophysiology studies will need to be conducted to determine how neuronal excitability of CGRPR^+^ cells changes as bladder pain persists over time.

In conclusion, our studies establish that CGRP-Rs contribute to the development of persistent bladder pain in a lateralized manner. One limitation of our study is that our experiments were not fully powered to detect sex differences. CGRP and CGRP-R antagonists and antibodies have been shown to display sex differences in many models of migraine^63,64^.

Additionally, while we have shown in our previous study that there are no differences in expression of CGRP receptors between the right and left^19^, we do not know if there are differences in CGRP receptor functionality across hemispheres. Therefore, future exploration of sex-differences and functionality of CGRP receptors will give more insight into the contribution of CGRP receptors to the development of persistent bladder pain.

## Author Contributions

LL, BP, AS, UC, PP, MK, AES, and AN contributed to data collection and analysis. LL, BP, and BK wrote the manuscript, and all authors reviewed the final manuscript. LL, BP, AS, IM, and BK contributed to experimental design.

## Acknowledgements

We thank the funding for this project from the National Institutes of Health grants F32DK128969 (LL), R56 AG084691-01A1 (IM), R01DK115478 (BJK), and The Burroughs Wellcome Fund BWF-1022337 (LL). Authors have no conflicts of interest to report.

