## Supplemental Files for "The role of amygdala calcitonin gene-related peptide receptors on the development of persistent bladder pain in mice"

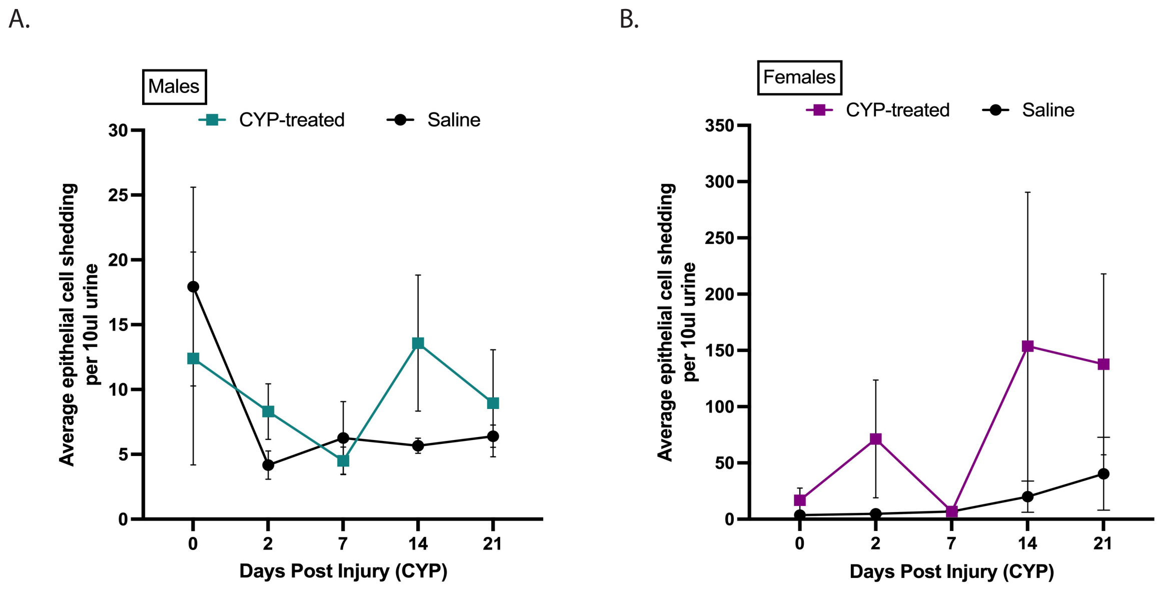


**S1: An increase in epithelial shedding was observed in female CYP-treated mice compared to males.** The average of epithelial cell shedding in the urine of **(A)** male and **(B)** female mice treated with either CYP (colored squares) or saline (filled circles). Data represent mean ± SEM. N=6/group.

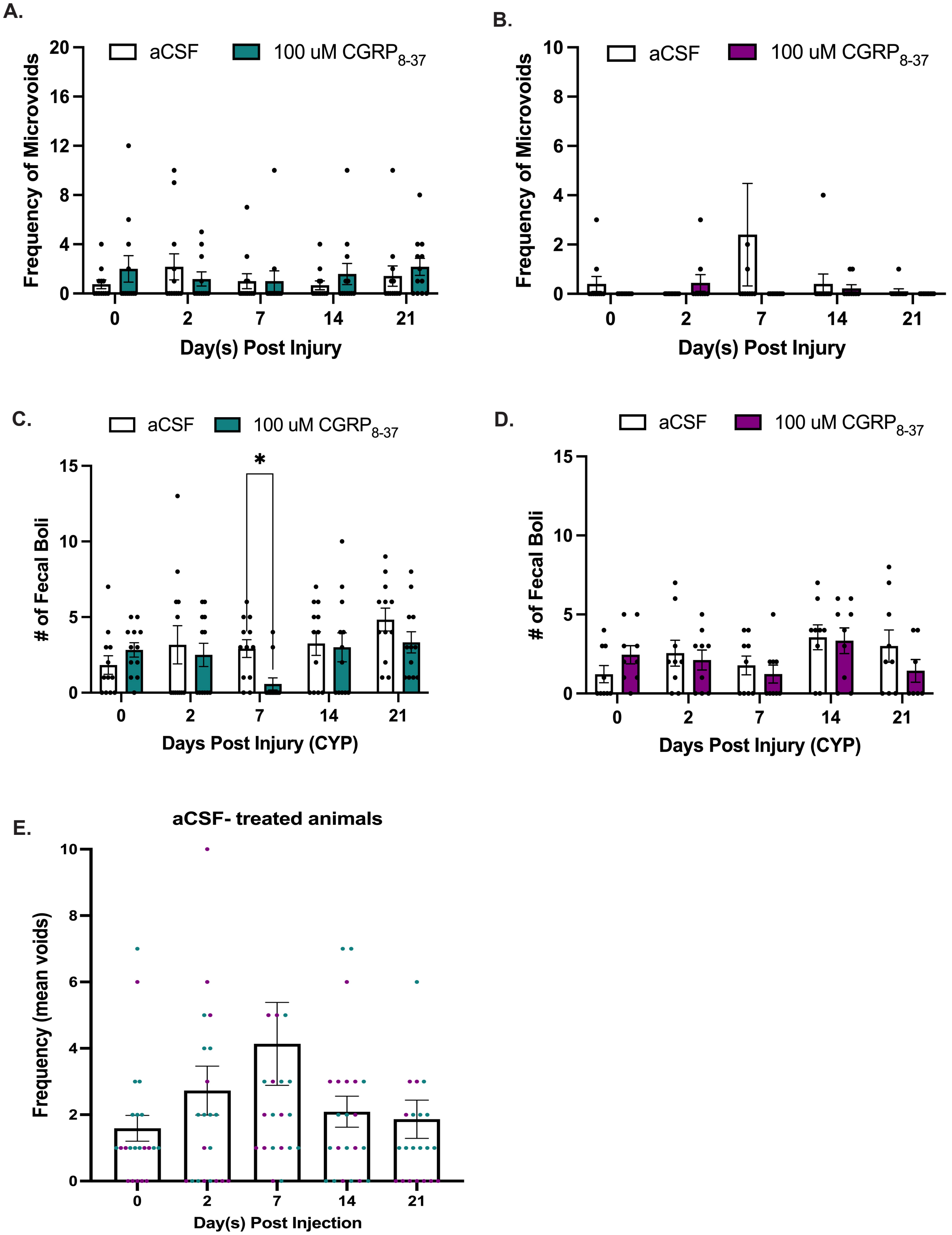
A.
Frequency of Microvoids
aCSF
100 uM CGRP8-37
Day(s) Post Injury
B.
Frequency of Microvoids
aCSF
100 uM CGRP8-37
Day(s) Post Injury
C.
### of Fecal Boli
aCSF
100 uM CGRP8-37
*
Days Post Injury (CYP)
D.
### of Fecal Boli
aCSF
100 uM CGRP8-37
Days Post Injury (CYP)
E.
aCSF- treated animals
Frequency (mean voids)
Day(s) Post Injection

**S2: CGRP$_{8-37}$ in the right or left CeA had no significant effect on the number of microvoids and minimal effect on fecal boli in CYP-treated mice. (A-B)** The frequency of microvoids was measured in CYP-treated animals that received either treatment of aCSF or CGRP$_{8-37}$ in the **(A)** left or **(B)** right CeA. The number of fecal boli was also measured in CYP-treated animals that received treatment in the **(C)** left or **(D)** right CeA. Šídák's multiple comparison's test revealed a significant difference in the number of fecal boli at 7 DPI when treatment was administered in the left CeA (*, $p < 0.05$). N=9-12/group. **(E)** The effect of CYP on voiding behavior in aCSF-treated mice. Purple dots represent animals that received aCSF in the right CeA, while teal dots represent animals that received aCSF in the left CeA.

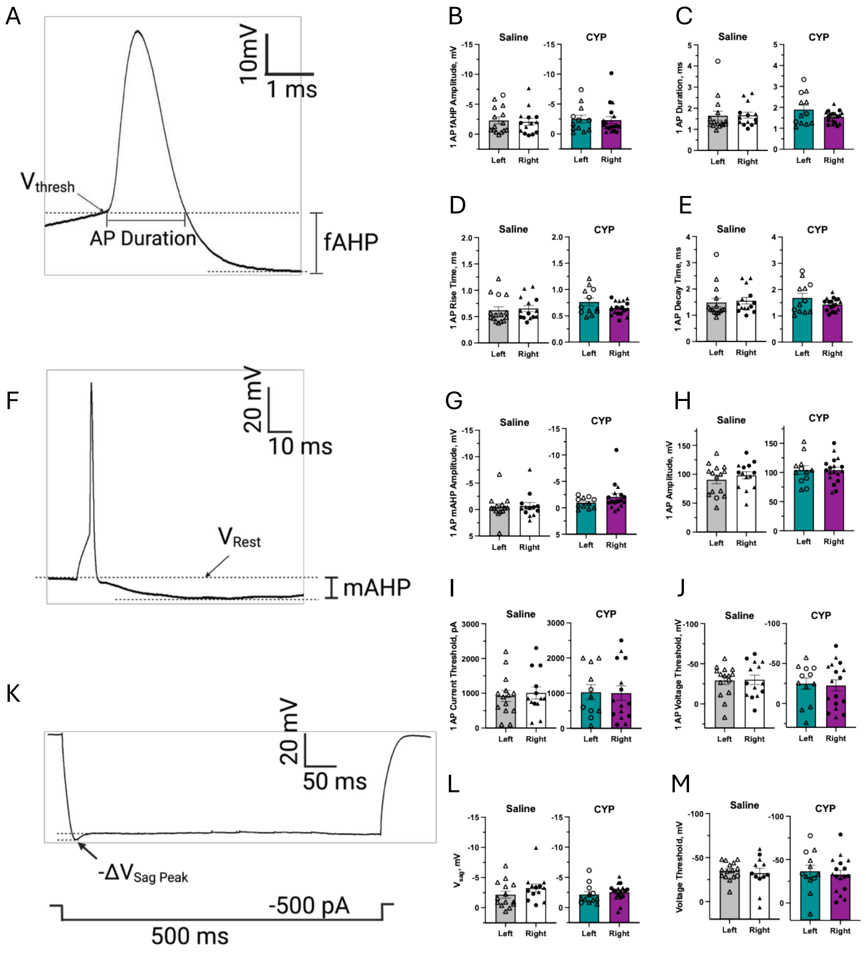
A
10mV
1 ms
Vthresh
AP Duration
fAHP
B
Saline
CYP
1 AP fAHP Amplitude, mV
Left
Right
C
1 AP Duration, ms
D
1 AP Rise Time, ms
E
1 AP Decay Time, ms
F
20 mV
10 ms
VRest
mAHP
G
1 AP mAHP Amplitude, mV
H
1 AP Amplitude, mV
I
1 AP Current Threshold, pA
J
1 AP Voltage Threshold, mV
K
20 mV
50 ms
-ΔVSag Peak
-500 pA
500 ms
L
Vsag, mV
M
Voltage Threshold, mV

**S3: No changes were observed in single action potential properties between hemispheres or CYP vs saline-treated animals**. **A.** Representative trace of an action potential. **B.** fast afterhyperpolarization peak from single action potential recording between left and right CeA in saline (left) and CYP group (right). **C.** Action potential duration of a single action potential between left and right CeA in saline (left) and CYP group (right). **D.** Single action potential rise time between left and right CeA in saline (left) and CYP group (right). **E.** Single action potential decay time between left and right CeA in saline (left) and CYP group (right). **F.** Representative trace of an action potential with a longer scale indicating the medium after hyperpolarization peak. **G.** medium afterhyperpolarization peak from single action potential recording between left and right CeA in saline (left) and CYP group (right). **H.** Single action potential peak amplitude between left and right CeA in saline (left) and CYP group (right). **I.** Current threshold to elicit a single action potential between left and right CeA in saline (left) and CYP group (right). **J.** Voltage threshold to elicit a single action potential between left and right CeA in saline (left) and CYP group (right). **K.** Representative voltage trace showing sag potential elicited with -500 pA, 500ms current step. **L.** Voltage sag peak values between left and right CeA in saline (left) and CYP group (right). **M.** Voltage threshold to elicit repetitive firing between left and right CeA in saline (left) and CYP group (right). All data are presented as mean $\pm$ SEM, and error bars represent SEM.**$p<0.01$, *$p<0.05$. Individual data points belonging to males and females are denoted with triangle and circle symbols, respectively. See Table1 for further statistical information.
